# Isolation of nuclei and downstream processing of cell-type-specific nuclei from micro-dissected mouse brain regions – techniques and caveats

**DOI:** 10.1101/2020.11.18.374223

**Authors:** M.C Chongtham, H Todorov, J.E. Wettschereck, S. Gerber, J. Winter

## Abstract

The mammalian brain consists of several structurally and functionally distinct regions equipped with an equally complex cell-type system. Due to its relevance in uncovering disease mechanisms, the study of cell-type-specific molecular signatures of different brain regions has increased. The rapid evolution of newer and cheaper sequencing techniques has also boosted the interest in cell-type-specific epigenetic studies. In fact, the nucleus holds most of the cell’s epigenetic information and is quite resistant to tissue dissociation processes as compared to cells. As such, nuclei are continually preferred over cells for epigenetic studies. However, the isolation of nuclei from cells is still a biochemically complex process, with every step affecting downstream results. Therefore, it is necessary to use protocols that fit the experimental design to yield nuclei of high quality and quantity. However, the current protocols are not suitable for nuclei isolation of small volumes of micro-dissected brain regions from individual mouse brains.

Additionally, the caveats associated with centrifugation steps of nuclei extraction and the effects of different buffers have not been thoroughly investigated. Therefore, in this study, we describe an iodixanol based density gradient ultracentrifugation protocol suitable for micro-dissected brain regions from individual mice using Arc^*creERT2 (TG/WT)*^.*R*26^*CAG-Sun1-sfGFP-Myc (M/WT or M/M)*^. This mouse model shows sfGFP expression (sfGFP*+*) in the nuclear membrane of specific stimulus activated cells, thereby providing a good basis for the study - nuclei isolation and separation of cell-type-specific nuclei. The study also introduces new tools for rapid visualization and assessment of quality and quantity of nascent extracted nuclei. These tools were then used to examine critical morphological features of nuclei derived from different centrifugation methods and the use of different buffers to uncover underlying effects. Finally, to obtain cell-type-specific nuclei (sfGFP*+* nuclei) from the isolated nuclei pool of high viscosity, an optimized protocol for fluorescence activated nuclei sorting (FANS) was established to speed up sorting. Additionally, we present a 1% PFA protocol for fixation of isolated nuclei for long term microscopic visualization.

## 1. Introduction

The mammalian mouse brain consists of a large number of complex cell types performing various functions. These cell types are equipped with an equally complex system of phenotypically and functionally different nuclei. Nuclei store the genetic and most of the sophisticated epigenetic information of a cell. They are also more resistant to mechanical assaults, thereby allowing unbiased representation of cell types - sensitive and insensitive to tissue dissociation processes (Tasic et al., 2018 as cited in Bakken et al., 2018; Krishnaswami et al., 2016). Therefore, with the recent advances in sequencing techniques, the interest in studying nuclei from functionally distinct micro-brain regions has increased rapidly.

Nuclei isolation is a complex process from a biochemical point of view (Graham, 2001). The process starts with chemical or mechanical disruption (Blobel and Potter, 1966) of the cytoplasmic membrane while retaining the integrity of the nuclear membrane. Extensive chemical or mechanical stress during any step of the isolation process can stimulate nuclear leakage, releasing chromosomal DNA. This can induce clumping of nuclei and thereby result in a decreased yield (Graham, 2002). Reduction in yield is particularly undesirable when rare biological materials of miniature size, for example, the micro-regions of the brain, are under investigation. Apart from the quantitative aspects, the nuclei can also suffer, qualitatively, from several unwanted biochemical effects during the isolation steps. One such step is “centrifugation” to separate the nuclei from the cell debris. The density-based ultracentrifugation technique reduces mechanical stress on the nuclei and is more widely used than the standard centrifugation technique. However, the choice of a proper gradient solution is essential for this step. For example, in the study by Mita and colleagues (2010), it was shown that Ficoll-based density gradients induced cytokine/chemokine production by islet cells as compared to iodixanol-based density gradients. In addition to these challenges, the presence or absence of “cushion layers” during the gradient ultracentrifugation could lead to significant differences in nuclei yield and quality as has already been shown for exosome isolation (Yamashita et al., 2016).

For nuclei isolation, such a study is yet to be conducted. Apart from these, nuclei isolation via a density-based ultracentrifugation technique is a time-consuming process. For studies using Accessible Transposable Assay of Chromatin (ATAC)-seq, Reduced Representation Bisulfite Sequencing (RRBS)-seq, Chromatin Immunoprecipitation (ChIP)–seq, rapid isolation of high-quality nuclei is essential. Currently, a large number of protocols for nuclei isolation exist. However, they are not tailored to micro-dissected tissues. Besides this, the pitfalls - proper choice of buffers, tissue dissociation methods, and reducing downstream processing duration - have not yet been appropriately addressed. Finally, different buffers used for collecting isolated nuclei can have a considerable impact on the quality and downstream processing. However, there is no conclusive data on this matter.

Therefore, our current study presents a compact nuclei isolation protocol tailored for micro-dissected single mouse brain regions, with a minimum volume limit of 1mm3. The yield and quality of such nuclei are determined using established as well as novel microscopy techniques introduced in this study. After establishing the nuclei isolation protocol for small brain regions (adaptation from Mo et al., 2015), we proceeded to study the effects of ultracentrifugation with and without a “cushion” layer on nuclei yield and quality. Certain protocol modifications were also added to reduce overall nuclei processing duration to suit the demand of short nuclei isolation protocols. We identified that buffers with different densities/compositions could affect the nuclei sorting process using Fluorescence Activated Nuclei Sorting (FANS). Besides this, the buffers can also affect nuclei integrity. Cumulatively, our study describes a protocol for isolating nuclei from small micro-dissected brain regions, tools and scripts for analyzing nuclei, downstream processing using FANS and appropriate choice of buffers for nuclei storage.

## 2. Results and discussion

### 2.1 Determining the nuclei yield efficiency of the selected protocol using whole brains

#### Choice of gradient solution and protocol

Amongst several existing nuclei isolation protocols for large tissue regions, we chose the method described by (Mo et al., 2015) as the starting basis for modifications to suit micro-dissected brain regions. The choice was based on the use of the iso-osmotic iodixanol as a gradient solution. As mentioned previously, several steps during the isolation process can lead to reduced quality and yield. One of them is the choice of gradient solution. Sucrose gradients have been the method of choice (Wilczok T and Chorazy K, 1960; Lovtrup-Rein and McEwen, 1966; Liao et al., 2020) for several organelles/cells for a long time. However, the use of sucrose gradients is gradually replaced by the less viscous, iso-osmotic iodixanol for the extraction of various cells/organelle/viral particles (Van Veldhoven et al., 1996; Graham, 2002; Cantin et al., 2008; Mita et al., 2010; Hutonorjs et al., 2012; Tauro et al., 2012; Katholnig et al., 2014; Marion-Poll et al., 2014; Mo et al., 2015; Onódi et al., 2018; Kovacovicova & Vinciguerra, 2019). This shift in trend is due to the better physiological resemblance of iodixanol solutions, leading to better preservation of cell organelles during the isolation step. Iodixanol-based density gradients have also been proven to be a better candidate than Ficoll-based density gradients (Mita et al., 2010; Quasem et al., 2017). For example, Quasem and colleagues showed the loss of “quiescence” in yeast cells when centrifuged with Ficoll-based density gradients compared to the former. Therefore, iodixanol is the gradient solution of choice.

#### Validation of the isolation process efficiency

For all our experiments, we used the mouse model Arc^*creERT2 (TG/WT)*^.*R*26^*CAG-Sun1-sfGFP-Myc (M/WT or M/M)*^ (see **Methods**). This mouse model shows nuclear membrane sfGFP expression in specific stimulus-activated cell populations, thereby providing a system for nuclei isolation experiments along with processing of specific nuclei (sfGFP+). Before the Mo et al., 2015, protocol could be adapted to smaller brain regions, it was necessary to determine the extraction efficiency. This was performed by using this protocol on adult whole mouse brains of the mouse line introduced above. This choice makes it easy for us to compare it to the percentage nuclei yield for adult brains available in the literature for Balb/cJ mice or rats. Though the lines are not exactly similar, we reasoned that, looking at nuclei yield percentages per organism would provide us with a good orientation of expected nuclei yield.

The tissue homogenization step is critical for nuclei yield. As such, we employed a douncing system (grinding pestle, see **Methods**), which can be readily applied to the tiny micro-dissected brain regions, instead of the douncers for large volumes of tissue, used by Mo and colleagues. The whole brain was micro-dissected into several pieces of a maximum of 5 cubic millimeters. 2-3 samples each were homogenized in the douncing setup and pooled before the centrifugation step (see **Methods**). Two independent nuclei isolation experiments from two biological replicates provided very similar amounts of nuclei (֊33×10^6^ nuclei per mouse, i.e., 30-40% of the whole brain tissue; nuclei counts were obtained using a hemocytometer). This number is in good agreement with the nuclei yield 80 x106 million/g of brain tissue reported in Yu et al., 2014 (30-40% of the adult mouse brain tissue, Balb/cJ). This amount is also higher than the ones reported by Sporn and colleagues,1962, (11% per rat brain) and Lovtrup rein and McEwen,1966 (20-25% per rat brain), where sucrose gradient solutions were used. Therefore, we concluded that this homogenization setup could be used for rapid and efficient nuclei isolation from small tissue regions of interest. **Figure 1A-D** provides a schematic representation of the nuclei isolation procedure as well as possible applications.

**Figure 1:**
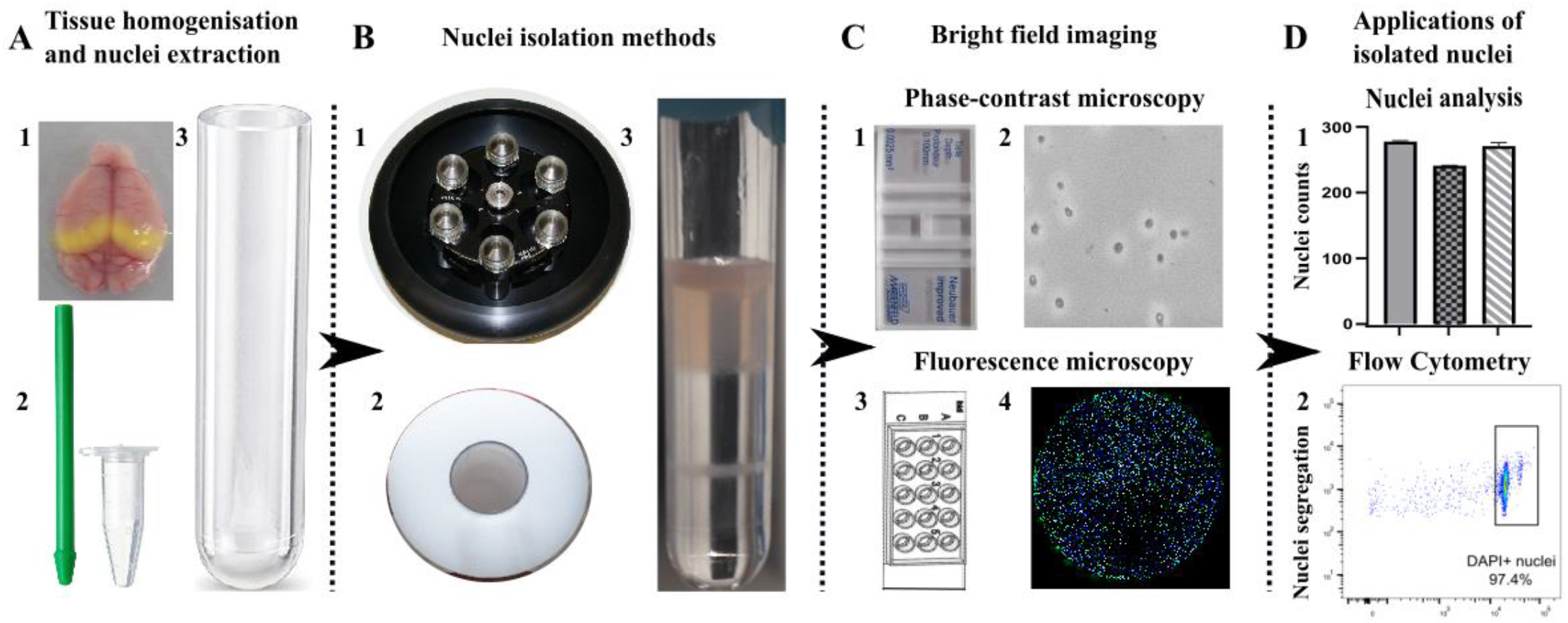
Schematic representation of cell-type specific nuclei isolation/processing along with possible applications: **A** Tools required for tissue homogenization of micro-dissected brain regions and gradient-based ultracentrifugation **1.** Whole brain tissue with hippocampus, highlighted in yellow **2.** A pestle to homogenize the micro-dissected tissue in a 1.5 mL reaction tube containing homogenization buffer **3.** 4 mL polypropylene ultracentrifugation tube to be used for layering the tissue homogenate over iodixanol gradients. **B** Equipments used for high speed ultracentrifugation **1.** HB6 ultracentrifuge rotor **2.** Polyvinylchloride (PVC) caste used to fit the centrifuge tube into the rotor tubes holders. **3.** Layer separation of the gradient iodixanol solution after ultracentrifugation of tissue homogenate **C** Methods for imaging nuclei isolated after the gradient-based ultracentrifugation. For phase-contrast microscopy, nuclei are loaded in a Neubauer chamber (**1**) and imaged using Leica DM IL inverted microscope. **2.** A representative image of nuclei under phase-contrast microscopy. For fluorescence microscopy, nuclei are loaded in a μ-slide angiogenesis chamber (IBIDI, **3**) and imaged using Leica AF7000 Widefield/SP5 confocal microscope. **4.** A representative whole-scan image of nuclei embedded in a single μ-slide chamber. **D** Downstream applications of extracted nuclei **1.** Representative image of data quantification of the nuclei obtained under different durations of ultracentrifugation using GraphPad Prism **2.** Segregation of desired nuclei using flow cytometry (BD FACSAriaSORP sorter).

### 2.2 Adaptation of the protocol to micro-dissected brain regions: Nuclei isolation and visualization

Upon verifying the nuclei yield, the protocol was modified to accommodate the small brain regions’ micro-volume requirements, including the hippocampus, hypothalamus, pituitary, pre-frontal cortex and nucleus accumbens (see **Methods**). The modifications would enable the optimum yield of nuclei from these brain regions from individual mice.

Tissues were dounce-homogenized in micro-volumes of supplemented homogenization buffer (see **Methods**). The obtained homogenates were loaded into 4 mL ultracentrifugation tubes containing microvolumes of iodixanol gradient solutions. Microvolumes were used to avoid nuclei dissipation, which could occur during ultracentrifugation, and nuclei collection using larger volumes. After 18 minutes of ultracentrifugation, nuclei were collected from the 30-40% gradient layer.

#### Nuclei visualization and yield determination

The nuclei, thus obtained, were initially visualized without trypan blue staining using phase contrast microscopy. Phase-contrast microscopy is one of the most straightforward techniques to determine nuclei yield and shape, which are critical to a successful isolation process. The yield was determined by manual counting of nuclei inside a hemocytometer using a Leica DM-IL inverted microscope (**Figure 2A-1**, n = 3 mouse replicates). The nuclei yield from the different brain regions with hippocampus > pituitary> PFC > hypothalamus > nucleus accumbens is summarized in **Figure 2A-2.** The small deviations from the average between replicates could be due to individual variabilities in micro-dissections of biological samples, douncing or nuclei collection from the 30-40% iodixanol solution layer. The representative images of the nuclei without trypan blue staining are shown in **Figure 2A-3** (nuclei of good quality appear darker). Though the nuclei yield can already be determined without trypan blue staining, it is hard to estimate nuclei integrity without the stain. Generally, a darker trypan blue stain indicates a more porous membrane (Zhu et al., 2016). This can be extended to indicate membrane integrity, a critical factor in determining the success of isolation. In our studies, we only compared nuclei from experiments where the staining patterns were similar. Representative images of trypan blue-stained nuclei from the different brain regions are shown in **Figure 2A-4**. Interestingly, trypan blue staining revealed small differences in the intensity patterns of nuclei coming from distinct areas. This could indicate different porosity levels of nuclear membranes within brain regions.

**Figure 2:**
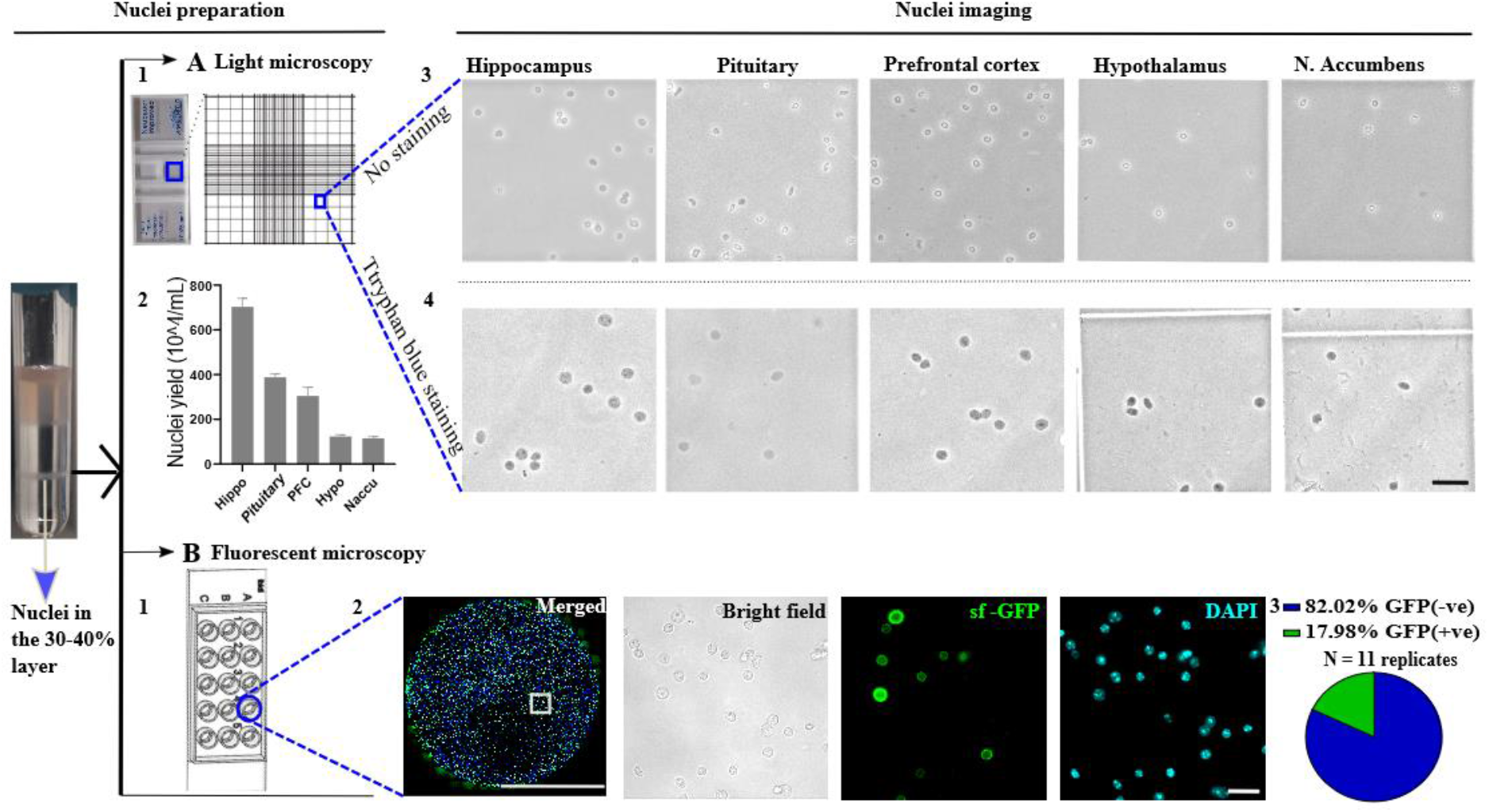
Data quantification and visualization using microscopy tools. **A**. Observation of isolated nuclei in a hemocytometer **(1)** using a Leica DM IL inverted microscope, 40x air objective. **2.** Average nuclei yield from 3 biological replicates in descending order of yield for hippocampus, pituitary, PFC, hypothalamus and nucleus accumbens. **3.** Representative images of nuclei from the respective brain regions (square area of 0.25mmx0.25 mm). **4.** Representative images from the same nuclei isolation but diluted with trypan blue in a ratio of 1:1. Scale bars correspond to 30 μm. Error bars represent +SEM **B.** Fluorescence microscopy images of collected nuclei in a preparation of a μ-angiogenesis slide IBIDI chamber **(1). 2.** Merged image of nuclei embedded in a well of the IBIDI chamber using, 40x water objective. Scale bar = 2 mm. The subsequent figures show individual images using different filters. Scale bar = 30 μm **3.** Manually calculated percentages of nuclei expressing sfGFP (GFP+ve) on the nuclear membrane as opposed to that without sfGFP expression (GFP-ve), calculated from n = 11 individual images.

Another tool for visualization is widefield fluorescence microscopy. Here, we introduce the use of IBIDI chambers for nuclei imaging. DAPI-stained nuclei were loaded in the wells of the μ-angiogenesis plate (IBIDI, **Figure 2B-1**) and observed under the Leica AF7000 widefield fluorescence microscope. Images are shown in **Figure 2B-2**. The use of the μ-angiogenesis plate removes the disadvantage of unfixed nuclei solutions drying out during the long image capturing process, prevalent on traditional slides. To our knowledge, this is the first study describing the use of IBIDI chambers to observe unfixed nuclei for short to extended periods with relative ease. This tool can be useful for experiments where estimations of fluorescent nuclei are required. The mouse line we used provided the perfect example as some “activated” nuclei (see **Methods**) express sfGFP on the nuclear membrane (sfGFP+ nuclei) while the non-activated show no sfGFP expression (sfGFP-). **Figure 2B-3** shows an example of manual counts of sfGFP+ nuclei from the total nuclei pool (n = 11 image replicates).

#### FIJI as a tool for nuclei analysis

In addition to the determination of yield and quality described above, the microscopic images can become a repository of additional information about the nuclei, given proper nuclei analysis tools. FIJI provides a user-friendly tool for the quantification of such information. While physical parameters of a single nucleus can be determined manually using the selection tools in FIJI (see **Methods**), multiple nuclei can be analyzed simultaneously using high throughput, semi-automated/ automated macroscript analyses (see **Methods and Supplementary Table S1**). **Figure 3A** shows a schematic overview of image analysis using FIJI. The analysis can be performed manually, where the nuclei to be analyzed are identified using the selection tools in FIJI. Alternatively, a semi-automated high throughput analysis can be done with a macroscript to detect nuclei. **Figure 3B** shows the percentage of sfGFP+ nuclei calculated using a semi-automated analysis on the same image set as in **Figure 2B-3.** The proportion of sfGFP+ nuclei shows a good agreement with the manual count. The slight difference in percentages between the manual and the semi-automated counts can be attributed to the difficulty in analyzing some clustered nuclei as separate entities even with the use of “watershed” function. However, this is not a big problem. In fact, with access to FIJI, a rapid evaluation of the extracted nuclei’s physical parameters, including their optical density and volumetric shape, can also be performed. The corresponding sfGFP+ detection by flow cytometry is also depicted in **Figure 3B,** showing a good correspondence to the microscopy results.

**Figure 3:**
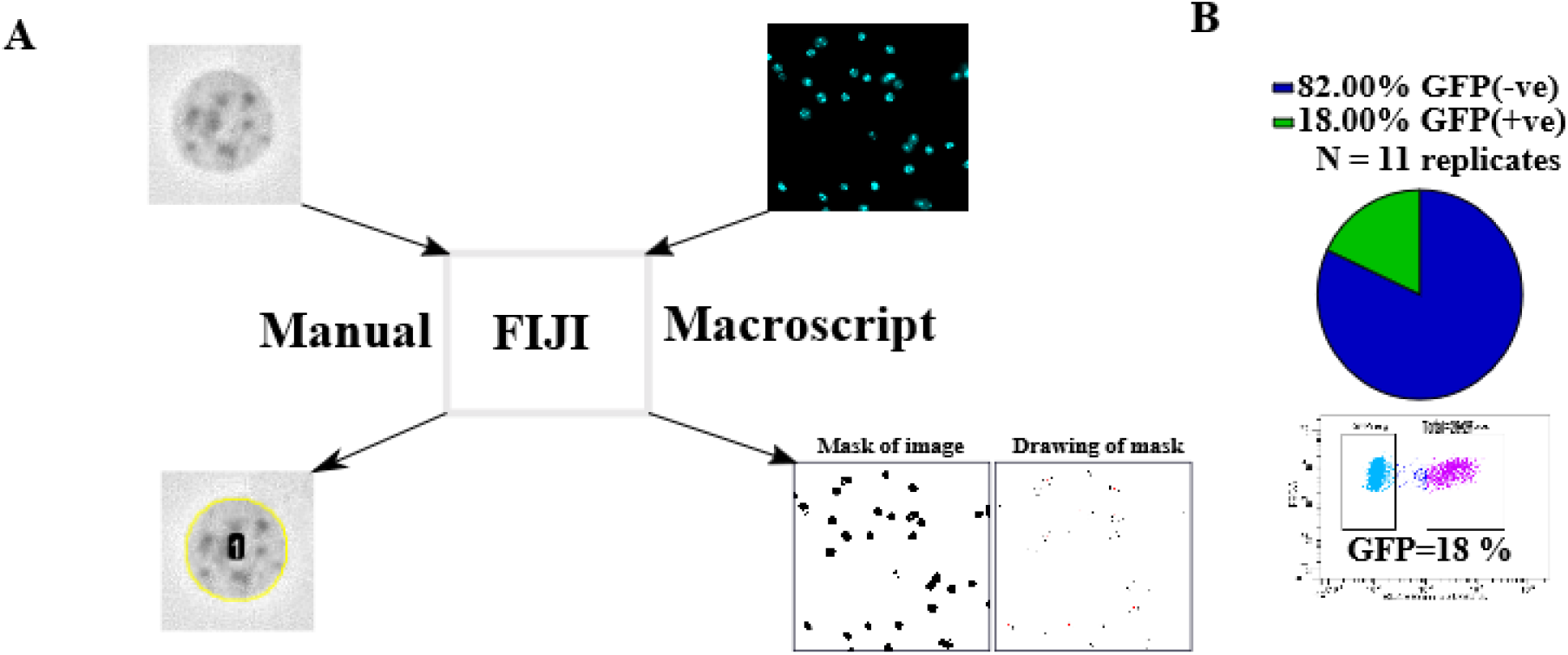
Image analysis using FIJI. **A.** Schematic diagram for the analysis of widefield microscopy images using FIJI. This can be performed either by selecting the region of interest manually or using macro to detect nuclei. **B.** shows a proof of principle for a macroscript to calculate the percentage of sfGFP+ nuclei (see **Methods**) n =11 images. The percentage of sfGFP+ nuclei detected by flow cytometry is shown below the graphical representation of the analyzed data.

In summary, we, successfully, applied this nuclei isolation technique to small brain regions including the nucleus accumbens. Therefore, it is assumed that this protocol can be extended to other small brain regions like the amygdala, dentate gyrus, ventral tegmental area, etc. To isolate nuclei from brain regions that are even smaller than those described here, e.g., the locus coeruleus, additional modifications not covered by this study need to be incorporated into the protocol.

Having established the nuclei isolation protocol, we next aimed at incorporating certain modifications to increase the nuclei yield and the speed of isolation. Furthermore, we tested different buffer compositions, as described in the following section.

### 2.3 Protocol modifications aimed at improving nuclei yield

A scheme of our current protocol with the “cushion” layer is shown in **Figure 4 1-4.** Nuclei isolated with the cushion layer demonstrated typical round/oval shape, characteristic of suitable nuclei quality (**Figure 4 3)**. The use of the cushion layer in this protocol, potentially, limits the mechanical stress experienced by the nuclei during the high-speed ultracentrifugation procedure. However, it is challenging to collect nuclei from the 30-40% cushion layer (indicated in **Figure 4 4**) with high consistency for such a small volume. This is partly due to the reduced visibility of accumulated nuclei at the 30-40% layer interface. During the nuclei collection from the thin 30-40% layer, pipette tip positions have to be continuously adjusted within the 30-40% layer to maximize nuclei collection. This could inadvertently lead to a loss of nuclei. Besides this, the high viscosity of the iodixanol gradient solution, making it unsuitable for sfGFP+ nuclei sorting using Fluorescence Activated Nuclei Sorting (FANS), where the sheath fluid usually is PBS. The underlying reason is that microfluidics’ laminar nature is necessary to adequately detect particles during flow cytometry (as mentioned in EP1242804A2, Thermo Fisher). However, this is disturbed when the viscosity contrast between adjacent laminas is large (Kurdzinski et al., 2017), as in our case. Like other studies using sucrose gradient centrifugation methods, before sorting, the nuclei have to be re-centrifuged and resuspended in a less viscous solution like PBS (Dammer et al., 2013).

**Figure 4:**
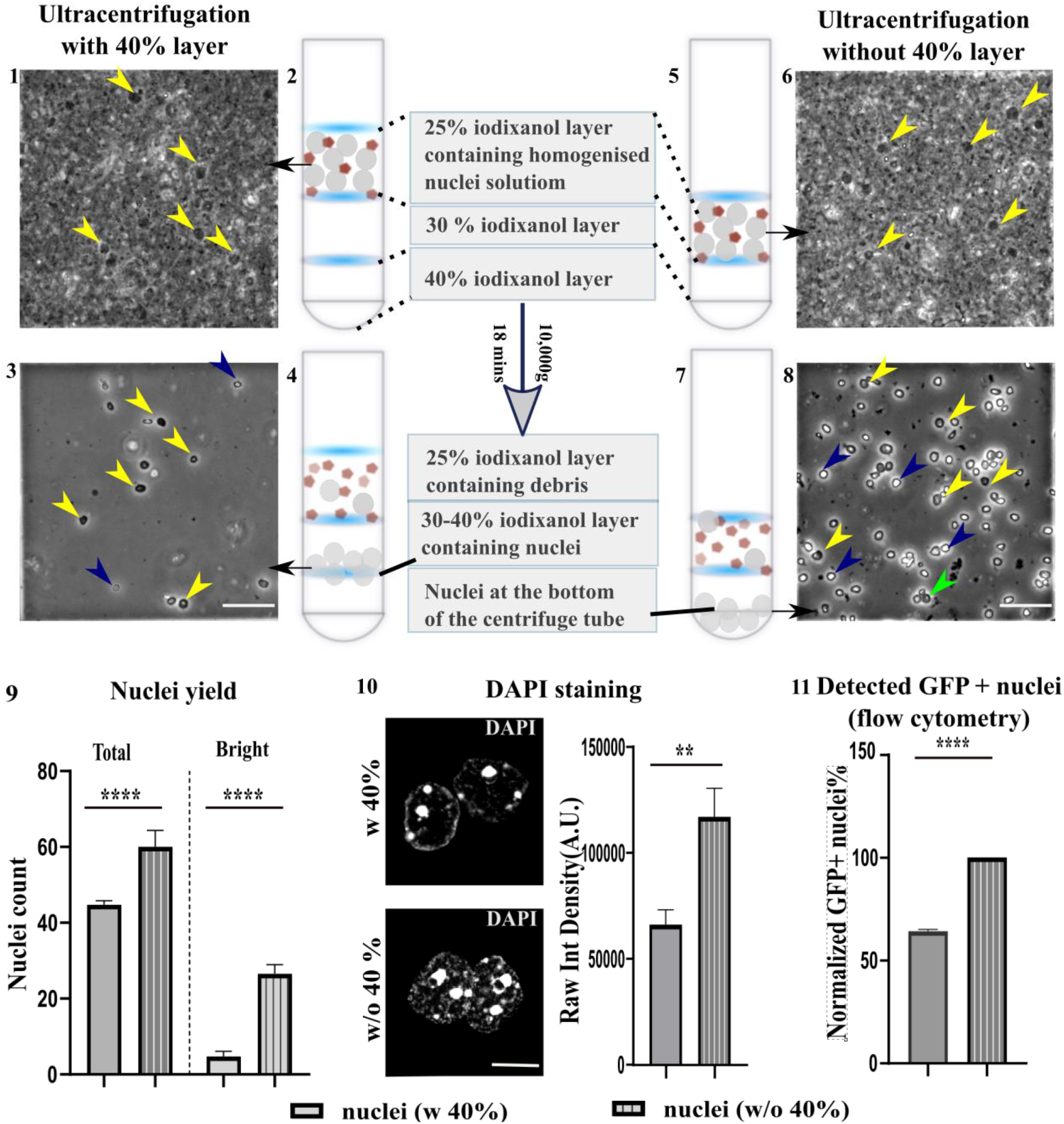
Comparison of ultracentrifugation with (w) and without (w/o) the 40% cushion layer. Schematic diagram of w 40% and w/o 40% is shown in 1-4 and 5-8, respectively. **1, 6** Phase contrast microscopy shows the tissue homogenate (nuclei and debris) in the 25% iodixanol layer before centrifugation in ultracentrifuge tubes (**2, 5**)**. 3, 8** Phase contrast microscopy images of nuclei collected from the 30-40% layer (**3**) or the pellet (**8**) after ultracentrifugation from tubes **4** and **7** as indicated in the figure. Yellow arrowheads indicate dark nuclei, blue arrows indicate the bright nuclei of smaller size and green indicates nuclei aggregation. Scale bar: 50 μm**. 9** Quantification of total nuclei yield along with bright nuclei obtained from the ultracentrifugation with and without the 40% layer is shown. Total nuclei count increased significantly (n=8, p<0.0001) in the centrifugation without the cushion layer. This was accompanied by a significantly higher number of smaller bright nuclei (n=8, p<0.0001). **10** Qualitative and quantitative differences (n = 7, p<0.01) in the DAPI staining pattern in the nuclei obtained from w 40% and w/o 40% layer (confocal microscope, 63x oil objective) Scale bars: 10 μm. **11** Normalized plots of percentage of sfGFP+ (GFP+) nuclei detected during sorting of nuclei obtained from w 40% or w/o 40% layer show significant differences in the percentage of detected GFP+ nuclei (n=5, p<0.0001). All statistical tests were performed using student’s t-test in GraphPad Prism. All error bars represent +SEM.

#### Effects of removing the cushion layer during ultracentrifugation

As an alternate experimental strategy to reduce the errors in collecting “floating” nuclei and increase yield, we attempted to remove the cushion layer during ultracentrifugation (**Figure 4 5-8**). This strategy also allowed direct resuspension of pelleted nuclei for FANS processing without the extra step of re-centrifugation. However, Yamashita snd colleagues (2016) reported in their study on exosomes that different types of centrifugation methods lead to different physicochemical properties. Meanwhile, Duong et al., 2019, demonstrated that the use of cushion layers increased the yield of extracellular vesicles. To our knowledge, despite the prevalence of several centrifugation modifications in brain nuclei extraction protocols, such a study has not been performed for nuclei isolation yet. Since our main aim was to obtain a high yield of good quality nuclei, we decided to examine the effects of removing the cushion layer as a starting point. We proceeded by evaluating the easily accessible and quantifiable parameters - nuclei yield, size, and optical density, which act as markers for efficient nuclei processing, using FIJI.

**Figure 5.**
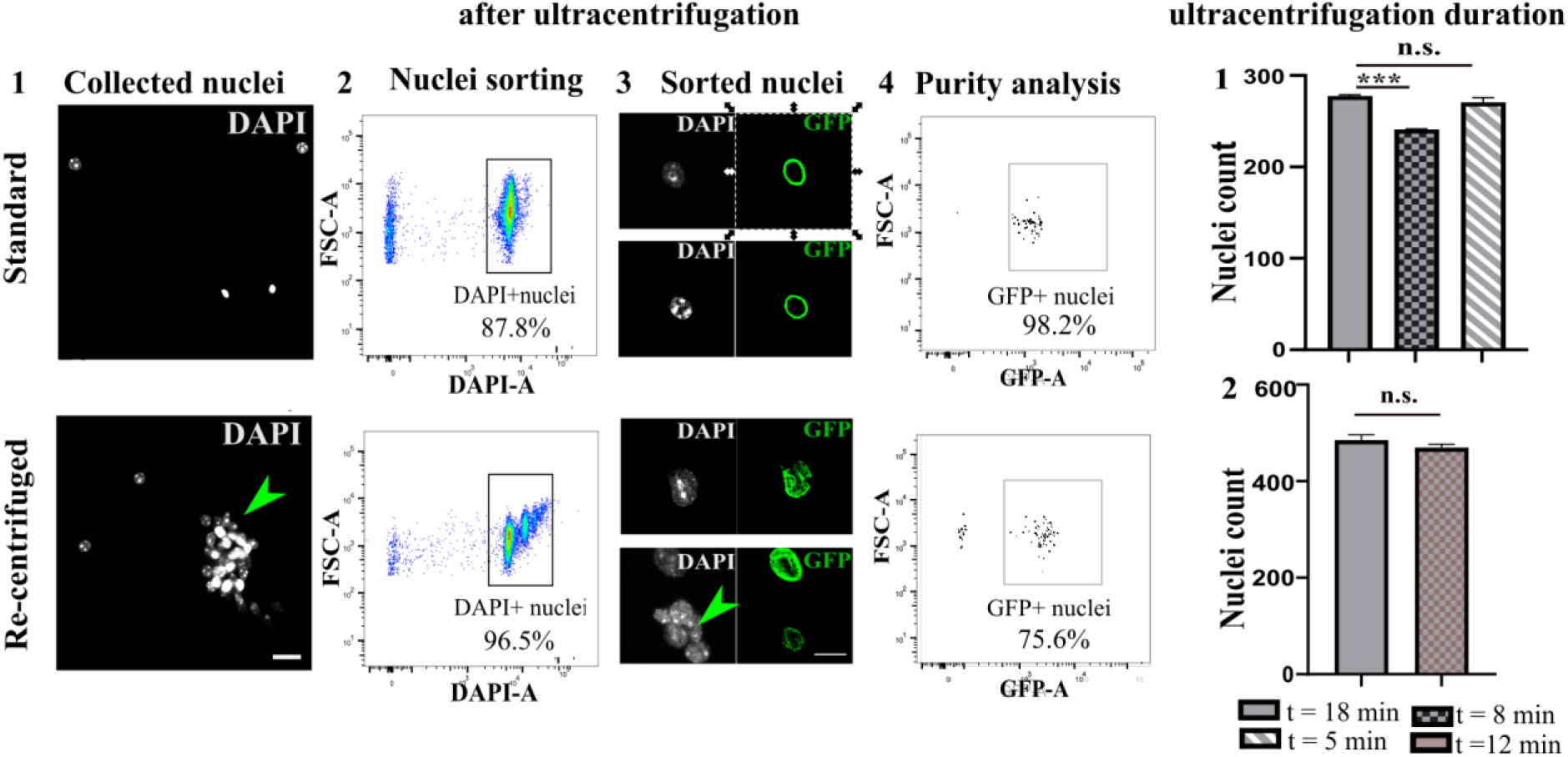
Methods to reduce nuclei processing duration. **A.** Differences between nuclei re-centrifuged with PBS/HB as compared to standard nuclei. **1.** Comparing microscopic images between normal and re-centrifuged nuclei reveals the occurrence of nuclei aggregates in re-centrifuged nuclei (indicated by green arrow). Scale bar: 30 μm. **2.** Dot plot of single nuclei from the corresponding centrifugation methods with the detection of aggregates in re-centrifuged nuclei. **3**. shows loss of integrity of sorted nuclei and the inclusion of sfGFP-nuclei cluster (green arrow) from re-centrifuged nuclei pool compared to standard nuclei. Scale bar: 10 μm. n = 3 replicates **4**. Reanalysis of the sorted nuclei detects a lower purity of re-centrifuged nuclei (representative image). **B.** Nuclei yield during different durations of centrifugation with the cushion layer. **1.** Graphical representation of differences in nuclei yield from the same homogenate when centrifuged for different durations ™18 min, 8 mins, and 5 minutes at 7820 rpm. **2.** Similar nuclei count between centrifugation for 12 and 18 minutes. n = 3 replicates each. All statistical tests were performed using a student’s t-test in GraphPad Prism. All error bars represent +SEM. (***, p<0.0005; n.s.- not significant).

Nuclei pellets obtained after ultracentrifugation of tissue homogenate **without** the cushion layer (w/o) were resuspended in homogenization buffer (HB/0.4% Igepal). These nuclei (w/o; **Figure 4 8**) were then compared to nuclei collected from the cushion layer from the homogenate ultracentrifuged **with**(w) the cushion layer (30-40% iodixanol layer, **Figure 4 4**). We observed a significant increase in nuclei yield under isolation performed w/o cushion layer over multiple experiments with a total analysis of 100-200 individual nuclei per large square (1mm2) of the hemocytometer (**Figure 4 9,** p<0.001, n= 8 replicates). However, this was accompanied by a significant increase in nuclei with a different refractive index than that extracted with the cushion layer (**Figure 4 9**). These nuclei appeared bright in the phase-contrast microscopy and, therefore, were termed as “bright nuclei”. The bright nuclei (indicated by blue arrows) were smaller and less opaque to light when compared to the normal nuclei (indicated by yellow arrows) **(Figure 4 4, 8**). The appearance of the bright nuclei cannot be entirely explained by the difference in the resuspension buffers (HB/0.4% Igepal vs. 30-40% iodixanol layer). This is because a high number of bright nuclei were still observed when a resuspension buffer of similar viscosity as the 30-40% iodixanol layer was used for dilution of the nuclei pellet (w/o cushion layer). Therefore, we concluded that ultracentrifugation without the cushion layer could technically lead to an increase in nuclei yield with some nuclei showing a different phenotype. Though we did not further investigate the origins of such nuclei, we speculate that the change in their refractive index could reflect the mechanical stress suffered by the nuclei during the pelleting. Concerning the nuclei yield, the ultracentrifugation with the cushion layer yields a lower quantity, perhaps due to potential loss of the nuclei by passive diffusion to the neighboring layers. In contrast, the pelleted nuclei have a reduced tendency of diffusion, and hence the nuclei yield is higher.

To further characterize the optical differences in the nuclei obtained from the two methods of centrifugation (resuspension of pellet from method w/o cushion layer with 30% iodixanol buffer), we used a TCS SP5 laser confocal microscope. We observed a difference in the staining pattern of DAPI in some of the nuclei extracted without the cushion layer. However, it was difficult to elucidate if these nuclei belonged to the group of “bright nuclei” from phase-contrast microscopy due to technical limitations. Using a 63x oil objective, we observed that DAPI staining had a more spotted pattern in the nuclei extracted without the cushion layer compared to the nuclei extracted with a cushion layer. The later showed concentrated staining of DAPI in specific regions **(Figure 4 10**). Nuclei with the highest observable differences were selected and analyzed using FIJI for the intensity of DAPI staining. Results in **Figure 4 10** showed that the total intensity was higher in the nuclei extracted without the cushion layer (student’s t-test, p<0.01, n=7). DAPI spots usually denote more regions of heterochromatin, and we suspect that the mechanical stress to which the nuclei were subjected to during pelleting, could technically lead to this observation. Interestingly, when these nuclei from w and w/o 40% layer (pellet resuspension with either HB-0.4% Igepal or 30% iodixanol) were sorted using FANS, sfGFP+ nuclei detection was significantly higher in nuclei pool extracted w/o 40% layer (**Figure 4 11**).

Conclusively, it is not advisable to perform ultracentrifugations without the cushion layer for sophisticated downstream epigenetic analyses, where the chromatin state should be well preserved as nuclei are sensitive to mechanical stress. The only advantage of ultracentrifugation w/o the cushion layer is the ease of resuspending pelleted nuclei with any buffer for isolating cell-type specific nuclei using FANS. For example, we observed that when using PBS or HB (both diluted with 0.4% Igepal, **see Methods**) as the resuspension buffer instead of 30% iodixanol, the sorting speed of sfGFP+ nuclei increased at least 10x as compared to using nuclei collected from the cushion layer as input for sfGFP+ nuclei sorting.

### 2.4 Protocol modifications aimed at increasing the speed of nuclei processing

As indicated previously, nuclei collected without the cushion layer can be easily resuspended in a buffer of choice for downstream processing, for example, separation of cell-type-specific nuclei using FANS. However, nuclei quality is reduced. Therefore, in agreement with the literature on extracellular vesicles (Duong et al., 2019), a cushion layer should be incorporated during nuclei collection. However, sorting with nuclei pool collected from the cushion layer is a rather slow process. Since the sorting speed is also critical for downstream analyses, we decided to re-centrifuge nuclei collected from the interface of the cushion layer with PBS (0.4% Igepal) or wash buffer (0.4% Igepal) and resuspend with the same solutions. In the following text, nuclei collected from the cushion layer, which did not undergo re-centrifugation, will be termed as “standard nuclei”.

#### Effects of re-centrifugation of nuclei collected from the cushion layer interface

For the process of re-centrifugation, standard nuclei were pelleted at a low speed (5000xg at 4°C) to reduce mechanical stress on the nuclei. The pellet was then resuspended using a buffer of choice: PBS (0.4% Igepal) or the wash buffer (0.4% Igepal). The speed of FANS increased by 10-fold compared to that of standard nuclei (in 30% iodixanol). However, there were also more nuclei clusters or aggregates, as observed in both the dot plot of the flow cytometer (despite proper gating) and fluorescence microscopy (**Figure 5A 1-2**). Clusters reduce good quality nuclei yield. Besides this, the occurrence of clusters is detrimental when low cell numbers from rare tissues are considered. Clustering could be due to improper resuspension methods after the pelleting or presence of damaged nuclei. Pelleting could lead to a higher amount of nuclei aggregation, as was shown for exosomes (Tauro et al., 2012; Jeppesen et al., 2014; Yamashita et al., 2016).

Examining the sorted nuclei under the fluorescence microscope, we observed that a small proportion of the nuclei sorted from the re-centrifugation method, lost their oval shape. Additionally, some sfGFP-nuclei were sorted together with the sfGFP+ nuclei as clustered nuclei. Representative images comparing nuclei sorted from either the re-centrifugation method or standard nuclei are shown in **FigurA 3.** Reanalysis of the sorted sfGFP+ nuclei from the two types of preparations by flow cytometry also indicated the presence of sfGFP+ nuclei compared to standard nuclei **(Figure 5A 4**). Therefore, re-centrifugation with any buffer is highly discouraged due to the associated time loss in re-centrifugation, unnecessary aggregation of nuclei, and higher chances of false-positive nuclei sorting from these aggregates although the sorting speed is increased. Since reducing the time of cell-type specific nuclei processing is highly essential for time-sensitive samples, we next looked at reducing the duration of ultracentrifugation with the cushion layer.

#### Effects of reducing the duration of gradient ultracentrifugation with a cushion layer

As observed, using a cushion layer reduces the chance of aggregation while simultaneously retaining the nuclear membrane integrity. As we use micro-volumes of gradient solutions for the micro-dissected brain regions, we hypothesized that less time is taken by a nucleus to reach the 30-40% gradient layer from the homogenate layer. This hypothesis follows from, t ∝ l, where t = time of centrifugation and l = sedimentation distance/ radial length during maximum centrifugation minus radial length during minimum centrifugation with the same centrifugal force (Stokes’ law; Livshits et al., 2015). Thus, we reduced the ultracentrifugation time from 18 minutes through 8 mins to 5 mins using tissue homogenates from neocortices. This was performed to determine the optimum time for centrifugation for such small volumes without compromising nuclei yield. As can be observed from the nuclei counts in **Figure 5B 1,** reducing the ultracentrifugation duration from 18 minutes to 8 minutes led to a significant nuclei loss. The unexpected higher yield of the 5 minutes centrifugation time could be attributed to the profuse nuclei that escaped the centrifugal force on the 25% layer. This could indicate an incomplete nuclei extraction from the tissue homogenate upon ultracentrifugation for 5 mins. Therefore, we chose another time point between 18 to 8 minutes to test our hypothesis on efficient time reduction during ultracentrifugation. As shown in **Figure 5B 2**, we observed that 12 minutes of ultracentrifugation yielded similar nuclei to that of 18 minutes. In terms of quality also, the 12-minute ultracentrifugation yielded similar results as observed by the experimenter. Therefore, this duration would be ideal for faster processing of multiple samples without significant nuclei loss and membrane integrity. The nuclei collected from the 30-40% layer can then be diluted with a solution of low surface tension (for example, wash buffer - HB with Igepal - or PBS) to speed up the nuclei sorting. This was indeed the case as shown in Fernandez-Albert et al., (2019), though at the time of performing our experiments, this study had not yet been published. However, different buffers can have various effects on the extracted nuclei in retaining nuclei integrity. Therefore, we assessed the impact of some standard buffers on nuclei integrity using easily quantifiable physical parameters.

### 2.5 Impact of different buffers on nuclei quality, detection, and storage

It is expected that buffers of different chemical composition will have a distinct impact on nuclei integrity in any step of the nuclei extraction process. However, this effect during downstream processing steps after nuclei isolation has not been investigated. In our experiments, we specifically looked at how different buffers influence nuclei integrity as well as nuclei sorting during flow cytometry.

#### Effects of different buffers on nuclei detection during flow cytometry

As demonstrated previously, nuclei sorting speed increased when re-centrifugation and resuspension with low viscosity buffer were performed. Another interesting observation is the increase in sfGFP+ nuclei detection in flow cytometry (**Figure 6A,** n= 7, p<0.0005) when nuclei were resuspended in buffers of low viscosity. We hypothesize that the increase in the sfGFP+ nuclei mostly results from a better detection in cytometry. This might be because of improper laminar flow in the instrument when there is a high viscosity contrast between neighboring laminar layers, for example, when nuclei collected from the 30-40% layer are used directly.

**Figure 6:**
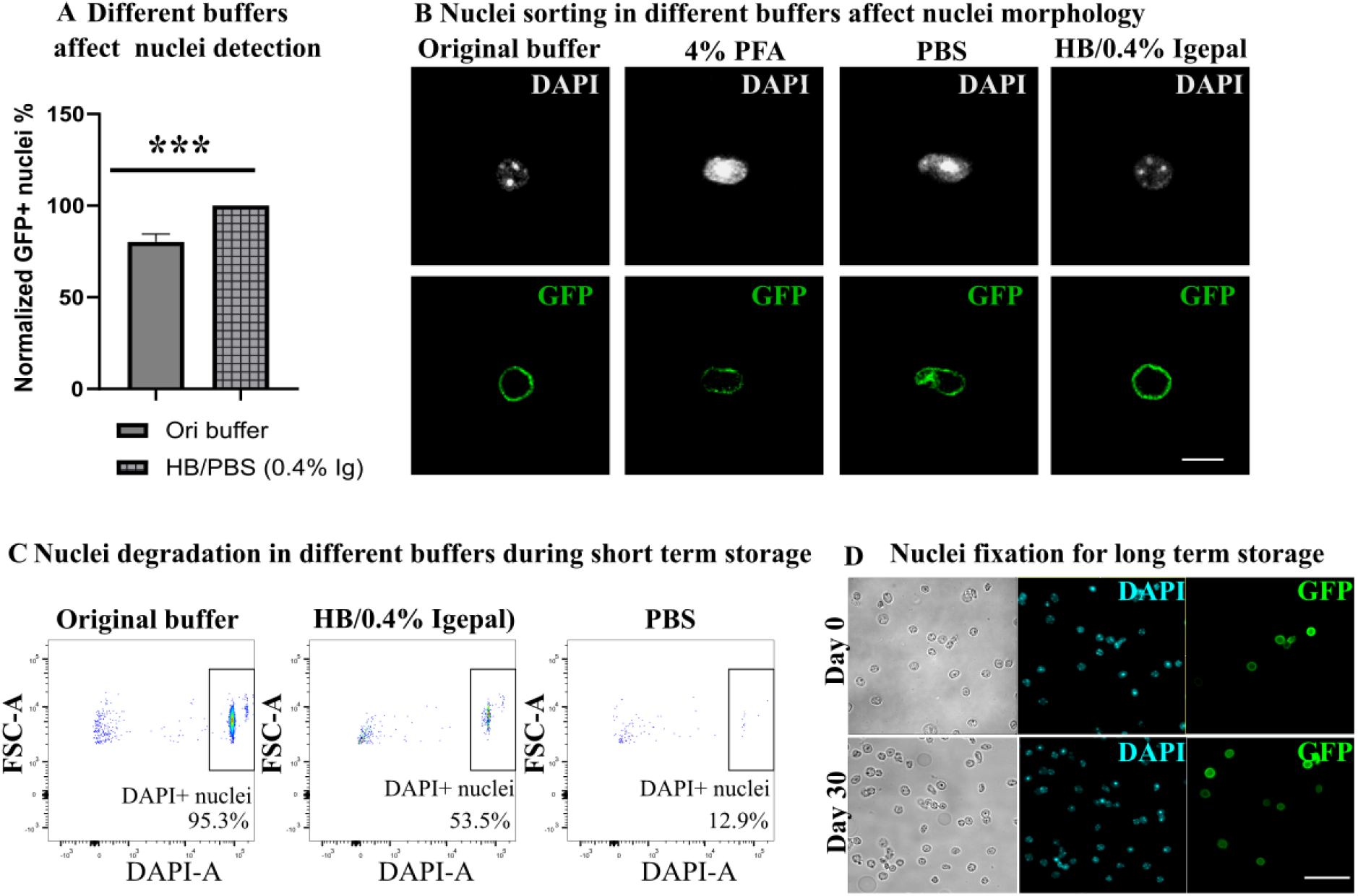
Effects of different buffers on nuclei detection and nuclei integrity: **A**. shows a significantly higher detection of sfGFP+ nuclei (GFP) when buffers of density lower than that of 30% iodixanol (0.4% Igepal, original buffer), i.e., PBS or HB, were used (n = 7, p<0.0005). **B**. Representative confocal microscopy images of sfGFP+ nuclei (GFP) sorted from the same nuclei pool into different buffers. Scale bar: 10 μm, Oil objective, 40x. **C.** Representative dot-plot of DAPI-positive nuclei in flow cytometry indicating the disintegration of nuclei stored in 30% iodixanol (0.4% Igepal) < HB (0.4% Igepal) < PBS over 24 hrs. **D.** Widefield microscopy images using different filters show preservation of 1% PFA fixed nuclei for 30 days as compared to the images from the day of nuclei isolation (Day 0). Scale bar: 50 μm, Water objective, 40x. All statistical tests were performed using student’s t-test in GraphPad Prism. All error bars represent +SEM.

#### Impact of different buffers on nuclei morphology

In order to examine the effects of different buffers on nuclei morphology, we decided to use “standard nuclei” to sort sfGFP+ nuclei into different collection buffers. This analysis revealed that nuclei sorted into HB (0.4% Igepal) had the closest nuclei integrity to that of 30% iodixanol diluted with HB (0.4% Igepal, 3:1 dilution hereby, termed as “original buffer”) as compared to PBS or PFA (**Figure 6B**). In fact, the sorted nuclei disintegrate when stored in PBS for more than a day (**Figure 6C**), while they remain morphologically stable in HB or original buffer for a week. Yamashita et al., 2016, also showed that the recovery rate of exosomes suspended in PBS was lower than when resuspended with BSA. However, the authors concluded that the lower recovery rate might be due to non-specific adsorption of exosomes to the cellulose during sterile filtration. Interestingly, Lima and colleagues (2018) reported the adverse effects of PBS in their study using iPSCs.

#### Nuclei storage for long term microscopic examination

We found 1% PFA to be the best reagent for the fixation of nuclei for long-term imaging. **Figure 6D** shows nuclei images before fixation and after a month of fixation in the IBIDI chambers and stored at 4°C. For long term preservation of nuclei, when necessary, we recommend storing the nuclei in a total volume of 70% glycerol in −80 °C (as in the protocol from www.collaslab.com).

## 3. Methods

### Animals and behavioral experiments

Male and female Arc^*creERT2 (TG/WT)*^.*R*26^*CAG-Sun1-sfGFP-Myc (M/WT or M/M)*^ (Denny et al., 2015; Mo et al., 2015) mice were used for the experiments. All mice were between 8 and 16 weeks old. For experiments where fluorescence microscopy was used, Tamoxifen (TAM − 150 mg/Kg, Sigma Aldrich; solvent- 1:9 of 100% ethanol: corn oil, Sigma Aldrich) was injected 5 hours before a behavioral stimulus. The stimulus is necessary to activate the Arc promoter leading to transcription of the CreERT2. In our case, the social interaction test (Krishnan et al., 2007) was used as the stimulus. TAM injection is necessary for the Cre protein to enter the nucleus and cleave the stop codon associated with the R26*CAG-Sun1-sfGFP-Myc (M/WT or M/M)*. This facilitates sfGFP expression on the nuclear membrane of cells activated by the stimulus. Mice were sacrificed 72 hours after TAM injection. All behavioral experiments were performed in accordance with the institutional animal welfare guidelines approved by the ethical committee of the state government of Rhineland-Palatinate, Germany (G-17-1-021)

### Nuclei isolation

#### Solutions

6x Tricine stock (pH-7.8, with KOH) was prepared with Tricine (120mM, Sigma, #SLBR4300V), KCl (150 mM, Roth), MgCl2 (30mM, Sigma, #011M0118V) and ddH2O and stored at 4°C. Stock solutions of spermine (150mM, Sigma, #BCBS6090V) spermidine (500mM, Sigma, #BCBW6017), cOmplete Mini EDTA-free protease inhibitor (1000x in ddH20, Roche, #29384100) were prepared in ddH2O and stored at −20°C for use within 2 months. 10% IGEPAL CA-630 (Sigma, 043K0654) stock solution was also prepared in ddH2O and stored at room temperature. All other solutions were prepared on the day of the experiments and can be stored for at least one day at 4°C

Homogenization buffer (HB) was prepared by dissolving sucrose (Sigma, #BCBT8436), resulting in a final concentration of 250mM in 1x Tricine stock solution (5 volumes of ddH20 and 1 volume of 6x Tricine stock). A small portion of this solution that was to be used for dounce homogenization was supplemented with a final concentration of 0.150 mM spermine, 0.500 mM spermidine, 1x cOmplete Mini EDTA-free protease inhibitor, and 1mM dithiothreitol (dTT, Applichem, #1P006802). The supplements were included to preserve nuclear membrane integrity and DNA better and to reduce protein degradation during the isolation process. This supplemented HB will be referred to as HB-supplemented in the following sections. The rest of the HB was used for preparing the different gradients from the original 60% iodixanol stock solution (Sigma, #BCCB9914). 50% iodixanol solution was prepared using five volumes of 60% iodixanol and one volume of 6XTricine stock. 40% and 30 % iodixanol solutions were subsequently prepared from the 50% iodixanol solution as recommended in “OptiPrep, Application Sheet C01” from “Alere Technologies” and stored on ice before use.

#### Protocol

The nuclei isolation protocol for neocortices (Mo et al., 2015) was modified following the requirements of micro-dissected tissues to increase nuclei isolation efficiency. All steps were processed at 4°C or on ice. The micro-dissected tissue samples were each transferred to 1.5 mL Eppendorf tubes containing 300 μL of supplemented HB. Tissues were dounced five times using the pestle (Z359971-1EA, Sigma Aldrich) in a soft twisting motion initially. 9 μl of 10% IGEPAL CA-630 was then added into each tube, achieving a final concentration of 0.3%. The addition of this non-ionic detergent is necessary to rupture the cytoplasmic membrane (Thoumine et al., 1999 and Caille et al., 2002 as cited in Tan et al., 2010) and the outer nuclear membrane. After that, the douncing was repeated eleven times with tight twisting rotations to get a good tissue homogenate with an orangish or whitish tinge depending on the initial amount of tissue (and RBCs) in the Eppendorf tube. An equal volume (300 μL) of 50% iodixanol was then added to this tissue homogenate, thereby bringing down iodixanol concentration to 25%. For routine experiments with the cushion layer, the prepared density gradients (40% iodixanol and 30% iodixanol) were layered as follows: 600 μL of 40% iodixanol followed by 600 μL of 30% iodixanol transferred gently on top of the 40% layer along the centrifuge wall with a pipette. The tissue homogenate in 25% iodixanol was then pipetted in a similar way to avoid disrupting the gradient formation. The tubes were then put in the white polyvinyl chloride caste to fit in the swinging bucket rotor (Sorvall HB-6) of the ultracentrifuge (Sorvall RC6+ Centrifuge). Ultracentrifugation was performed at 7820 rpm (10,000x g) at 4° C for 18 mins for all experiments except for studying different centrifugations times. Nuclei were collected (300 μL) from the 30%-40% iodixanol layer interface in a slow swirling motion with a 1000 mL pipette and transferred into a 1.5 mL Eppendorf tube through a 20 μm strainer (Partec 04-0042-2315). Thus, the collected solution is termed as “nuclei solution”. For flow cytometry, the nuclei solution was diluted with HB/0.4% Igepal) at a ratio of 3:1. This solution is referred to as the original buffer (OB).

The following modifications were made to determine the effects of the various types of centrifugation:

1. For validation of nuclei extraction efficiency performed by using the whole brain, the brain regions were micro-dissected into smaller pieces. 2-3 pieces of the micro-dissected tissues were independently homogenized in 1.5 mL Eppendorf tubes containing the HB. The homogenization was performed with the pestle (Z359971-1EA, Sigma Aldrich), which can be readily applied to the tiny micro-dissected brain regions instead of conventional tools for large tissues. The homogenates were then pooled to give a total of 5mL. The pooling was performed as we were interested in determining the nuclei yield from the whole brain, and separate ultracentrifugation for each homogenate could contribute to nuclei loss. This would not correctly represent the effective nuclei yield when the whole brain tissue was considered The collected solution was then diluted with an equal volume of 50% iodixanol to give 10mL of 25% iodixanol-homogenate mixture. The 50mL ultracentrifuge tube (FisherScientific, #12704868) was then layered with equal volumes of 40% and 30% iodixanol solutions. The 25% iodixanol-homogenate layer was then pipetted on top of the 30% iodixanol layer.
2. For Nucleus accumbens (roughly one cubic mm), douncing was performed using a 0.1 mL tissue grinder (Art. No. 0296.1, Roth) with less than 100 μL homogenization buffer. Igepal concentration and douncing repetitions were applied as in the above sections. After the initial douncing, volume was brought up to 150 μL by adding a homogenization buffer. The homogenate was mixed with equal volumes of 50% iodixanol and centrifuged similarly to other brain regions with gradients 40% (500 μL) and 30% (500 μL). 150 μL of nuclei were collected from the 30-40% nuclei layer.
3. For the experiments involving comparison of “with and without” cushion layer, 800 μL was used for each of the 40% and 30% gradients instead of 600 μL. The volume change was, specifically, made to minimize damage to the nuclei during pelleting without the cushion layer. After ultracentrifugation, for the experiments without the 40% layer, the upper homogenate layer and 30% layer were removed entirely using a 1000 mL pipette, leaving only about 50 μL at the bottom of the tube. The remaining nuclei solution was resuspended with either 30% iodixanol: HB with 0.4% Igepal (3:1) or HB (with 0.4% Igepal) or PBS with 0.4% Igepal for various purpose.
4. For nuclei count comparisons from different ultracentrifugation times, ultracentrifugation was performed for 12 mins, 8 mins, and 5 mins. The order of centrifugation with the set time was alternated between repeated replicate experiments to avoid unwanted errors introduced by the storage of isolated nuclei on ice for the length of the experiment per day.
5. For re-centrifugation, equal volumes of PBS or HB were added to the nuclei solution collected from the cushion layer and centrifuged at 5000x g. The supernatant was removed, and the pellet was resuspended with PBS or HB.

### Flow cytometry and fluorescent-activated nuclei sorting (FANS)

Flow cytometry analysis and FANS were performed using a BD FACSAria III SORP equipped with four lasers (405 nm, 488 nm, 561 nm, and 640 nm) and a 70 μm nozzle. GFP expression was detected using the blue laser and a 530/30 BP filter, whereas DAPI was detected using the violet laser and a 450/50 BP filter. Prior sort, 10,000 total events were recorded and a gating strategy was applied: first, nuclei were gated according to their forward- and side- scatter properties (FSC-A/SSC-A), followed by doublet exclusion using SSC-A and SSC-W. Nuclei were then gated according to their DAPI expression. GFP expression was used as a sorting gate. Sorted nuclei were collected in 1.5 mL Eppendorfs in PBS (with 0.4% Igepal), HB (with 0.4% Igepal), 30% iodixanol: HB with 0.4% Igepal (3:1) or 4% PFA to determine the effects of different buffers on nuclei integrity. The analysis was done using the BD FACSDiva 8.0.2 Software or FlowJo (v.10.6 or higher). Reanalysis of the sorted nuclei was performed by determining the percentage of sfGFP+ nuclei from at least 50 single DAPI positive nuclei.

### Microscopy

#### Phase-contrast Microscopy

10 μL of the nuclei solution was pipetted on a hemocytometer (Neubauer) for visualization under the (Leica DM IL inverted) microscope. For experiments involving trypan blue staining, the nuclei solution was diluted with trypan blue at a 1:1 ratio. Images were captured using 20x/0.3, air objectives, or 40x/0.5, air objective. Nuclei counting was performed in the usual format for hemocytometers – an average of the nuclei count from the four 1 mm by 1 mm squares multiplied by 10k/mL. For the nuclei counts comparing bright vs. total nuclei in **Figure 4 9**, counts from each of the 1 mm by 1 mm square, were considered a technical replicate.

#### Wide field Fluorescence Microscopy

The wells in the μ-slide angiogenesis plate (IBIDI, Cat.No: 81506) were incubated with a layer of polyethylenimine (PEI, 20 μL) for 20 mins and then washed with 1x PBS and HB (with 0.4% Igepal) subsequently. 10 μL of the nuclei solution was then loaded into the inner well of the chamber gently and incubated for 5 to 10 minutes. Nuclei were then visualized using the Leica AF7000 Widefield microscope with 40x/1.1 water objective using bright field, DAPI filter (A4), and GFP filter (L5). Tile scans were stitched using in-built software in the Leica LASX system.

Nuclei embedded in the PEI were fixed using 1% PFA after removing supernatant solution from the IBIDI wells for long-term imaging. After an incubation period of 5 minutes, the wells were washed with 15 μL of HB (with 0.4% Igepal) twice for 5 minutes each. A final volume of 10 μL of HB (with 0.4% Igepal) was pipetted into each well. The chamber was tightly sealed and stored at 4 °C for a month.

#### Confocal Microscopy

Leica TCS SP5 laser confocal microscope was used for all confocal microscopy images.10 μL of nuclei solution was pipetted on “SuperFrost Plus” microscopic slides from ThermoFisher Scientific. The solution was then covered with a round coverslip with a thickness of 0.17 mm and diameter 1.5 cm for viewing the nuclei through the inverted objective lens. Images were captured using 20x/0.7, air objective, 40x/1.1 (oil objective immersion) and 63x/1.4 (oil objective). For storing the samples for a few days, the coverslips were sealed with commercially available nail polish.

### Image analyses

All microscopic images were analyzed using Fiji.

A. For manual analysis of single nuclei, the steps are shown as hereunder:

1. Click “File”
2. Open “Image”
3. Click “Process” -> “Subtract background”
4. Click “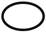” -> right-click and select “elliptical selections”
5. Draw “ellipse” around the nuclei of interest manually
6. Go to “Analyze”->“ Set Measurements”-> select the required parameters from the menu and press “ok”
7. Go to “Analyze” -> “Measure”
B. For semi-automatic analysis of images, the following macroscript was used for green (I) and for blue (II) channels respectively for Section 3 of the results.

~~~
(I)
//selecting images
run(“Duplicate…”, “title=green duplicate channels=2”);
run(“Gaussian Blur…”, “sigma=1”);
//setting threshold
setAutoThreshold(“Intermodes”);
//run(“Threshold…”);
setThreshold(187, 1077);
setOption(“BlackBackground”, false);
run(“Convert to Mask”);
run(“Fill Holes”);
run(“Watershed”);
//measurements
run(“Set Measurements…”, “area mean min perimeter fit shape feret’s integrated display add
redirect=None decimal=3”);
run(“Analyze Particles…”, “size=15.00-500.00 show=Outlines display exclude”);
call(“ij.plugin.filter.ParticleAnalyzer.setFontSize”, 25);
(II)
//selecting images
run(“Duplicate…”, “title=blue duplicate channels=3”);
run(“Gaussian Blur…”, “sigma=1”);
//setting threshold
setAutoThreshold(“Intermodes”);
//run(“Threshold…”);
setThreshold(760, 65535);
setOption(“BlackBackground”, false);
run(“Convert to Mask”);
run(“Fill Holes”);
run(“Watershed”);
//measurements
run(“Set Measurements…”, “perimeter shape feret’s integrated display add redirect=None decimal=3”);
run(“Analyze Particles…”, “size=15.00-500.00 show=Outlines display exclude”);
//display characteristics
call(“ij.plugin.filter.ParticleAnalyzer.setFontSize”, 25);
~~~

For automatic analysis of multiple files, see **Supplementary Table S1**. For determining intensity of GFP signals, see **Supplementary Table S2**.

### Statistics

All statistical calculations were performed using GraphPad Prism (8.4.2). Student’s unpaired t-test with the assumption that the SD between populations is equal was used for statistical comparisons of two groups. In all calculations, p<0.01 was considered statistically significant, in order to decrease chances of false positives.

## Acknowledgement

We would like to thank the members of the IMB core facilities of microscopy and flow cytometry for helping us complete this paper. We appreciate the efforts of Maria Hanulova in helping us to find techniques to visualise unfixed nuclei and Stefanie Moeckel for flow cytometry. We also extend our sincere thanks to the Mouse Behavioral Unit of the Leibniz Institute of Resilience Research.

## Author contributions

MCC designed the project and performed the experiments. MCC and HT drafted the paper. JEW performed part of the experiments. SG edited the paper. JW supervised the project and edited the paper

## Notes

### Competing Interest Statement

The authors have declared no competing interest.

