## Supplementary tables for "Isolation of nuclei and downstream processing of cell-type-specific nuclei from micro-dissected mouse brain regions – techniques and caveats"

**Table S1: Macroscript for automatic analysis of nuclei images**

```
dir1 = getDirectory("Choose a Directory");
dir2 = getDirectory("Choose a Directory");
list = getFileList(dir1);
//run("Bio-Formats Macro Extensions");
	for(i = 0; i < list.length; i++){
		if(endsWith(list[i], ".tif")){
			filepath = dir1 + list[i];
			open(filepath);
			run("Duplicate...", " ");
			run("Gaussian Blur...", "sigma=1");
			setAutoThreshold("Intermodes");
			//run("Threshold...");
			setThreshold(1600, 65535);
			setOption("BlackBackground", false);
			run("Convert to Mask");
			run("Fill Holes");
			run("Watershed");
			run("Set Measurements...", "area mean min perimeter fit shape feret's
integrated display add redirect=None decimal=3");
			run("Analyze Particles...", "size=15.00-500.00  show=Outlines display");
			call("ij.plugin.filter.ParticleAnalyzer.setFontSize", 25);}
		maskName = replace(list[i], ".tif", "_mask.tif");
			saveAs("Tiff", dir2 + maskName);
			drawingName = replace(list[i], ".tif", "_drawing.tif");
			saveAs("Tiff", dir2 + drawingName);}
saveAs("Results", dir2 + "Results.csv");
run("Close All")
```

**Table S2: Macroscript for measuring intensity of sfGFP fluorescence in the nuclear membrane**

```
roiManager("reset");
run("Clear Results");
name=getTitle();
run("Duplicate...", "title=blue duplicate channels=3");
setAutoThreshold("Otsu dark");
run("Convert to Mask");
run("Erode");
run("Erode");
run("Erode");
```

```
run("Erode");
run("Analyze Particles...", "size=7.00-500.00 show=Outlines add");
run("Set Measurements...", "mean redirect=None decimal=3");
selectWindow(name);
run("Duplicate...", "title=green duplicate channels=2");
for (i=0;i<roiManager("count");i++) {
	roiManager("select",i);
	run("Make Band...", "band=1");
	run("Measure");
}
```
